# Comparative brain structure and the neural network features of cuttlefish and squid

**DOI:** 10.1101/2022.05.08.491098

**Authors:** Wen-Sung Chung, Alejandra L. Galan, Nyoman D. Kurniawan, N. Justin Marshall

## Abstract

Cuttlefishes, like their octopus cousins, are masters of camouflage by control of body pattern and skin texture to blend in with their surroundings for prey ambush and threat avoidance. Aside from significant progress on the cuttlefish visual perception and communication, a growing number of studies have focused on their behavioural neurobiology and the remarkably rapid and apparently cognitively complex reactions to novel challenges such as spatial learning to solve maze tasks and vertebrate-like cognitive capabilities (e.g. object recognition, number sense and episodic-like memory). Despite intense interest of cuttlefish, much of our knowledge of its neuroanatomy and links to behaviour and ecology comes from one temperate species, the European common cuttlefish, Sepia officinalis. Here we present the first detailed comparison of neuroanatomical features between the tropical cuttlefish and squid and describe differences in basic brain and wiring anatomy using MRI-based techniques and conventional histology. Furthermore, comparisons amongst nocturnal and diurnal cuttlefish species suggest that the characteristic neuroanatomical features infer interspecific variation in visual capabilities, the importance of vision relative to the less utilised chemosensory system and clear links with life modes (e.g. diurnal vs nocturnal), ecological factors (e.g. living depth and ambient light condition) as well as to an extent, phylogeny. These findings link brain heterogeneity to ecological niches and lifestyle, feeding hypotheses around evolutionary history and provide a timely, new technology update to older literature.

## Introduction

Cuttlefish, squid and octopus are the three groups of coleoid cephalopods exhibiting diverse adaptations in body form, life modes and behavioural repertoires. This is reflected in the underlying nervous system (Nixon and Young, 2003, Hanlon and Messenger, 2018). While the fourth extant group of cephalopods, Nautilus, has an obvious external shell, gas-filled and used for floatation, octopus and squid have lost almost all remnants of this ancient feature and may therefore inhabit a broad range of ocean depths (0-6000 m) (Jereb and Roper, 2010, Jereb et al., 2014). The cuttlefish possess an internal chambered cuttlebone that, while giving internal strength, is also controllably gas-filled and therefore the risk of implosion limits their living depth to above 400m. Interestingly, for unknown reasons, they also have a limited geographic distribution (high diversity in the Indo-Pacific but absence in the Americas and polar regions) (Sherrard, 2000, Jereb and Roper, 2005, Lu and Chung, 2017). The cuttlebone allows buoyancy control by adjustment of the ratio between air and liquid and cuttlefish can therefore hover in the water column or bury themselves in sand to hide. This hovering, usually close to the benthos, is in contrast to the continual swimming activity of squid and the almost exclusively benthic existence of coastal octopus (Denton and Gilpin-Brown, 1961, Hanlon and Messenger, 2018).

A growing number of *in situ* observations of cuttlefish species show that they are not solitary, as are most of the species of neritic octopuses and also not as social as schooling species of squid (Hanlon and Messenger, 2018, Lu and Chung, 2017). Cuttlefish may therefore have a partially social life and are known to aggregate, sometimes in large numbers, for breeding on a seasonal basis (e.g. European common cuttlefish, *Sepia officinalis*; Broadclub cuttlefish, *Sepia latimanus*; Australian giant cuttlefish, *Sepia apama*) (Norman et al., 1999, Hanlon et al., 2005, Yasumuro et al., 2015, Drerup and Cooke, 2021).

Cuttlefishes, like their octopus cousins, are masters of camouflage by control of body pattern and texture to blend in with their surroundings and use this ability both for prey ambush and threat avoidance (Marshall and Messenger, 1996, Chiao and Hanlon, 2001, Hanlon and Messenger, 2018, Gonzalez-Bellido et al., 2018, Osorio et al., 2022). In fact, cuttlefish spend most of their time in very effective and totally colourblind camouflage, they may also rapidly switch colouration to emphasise their presence, produce startle threats, attract mates or indeed cheat rival males (Norman et al., 1999, Hanlon et al., 2005, Zylinski et al., 2011, Brown et al., 2012, Chung and Marshall, 2016, How et al., 2017, Alejandra et al., 2020). The ability to alter their visual appearance is driven by neurally-controlled chromatophore (colours) and muscular hydrostat (papillae) systems coordinated by a set of brain lobes organised hierarchically (e.g. the simplest circuit, optic lobe (OPL) - lateral basal lobe (lB)– chromatophore lobe (Ch)) (Messenger, 2001, Gonzalez-Bellido et al., 2018).

The cuttlefish central nervous system (CNS) is built around a circum-oesophageal set of lobes that have expanded greatly, in part in response to their complex visual system and rapid visual-motor reactions (i.e. ballistic tentacular strike, visual communication) (Tompsett, 1939, Sanders and Young, 1940, Boycott, 1961, Messenger, 1968, Chichery and Chichery, 1987). The general shape of the cuttlefish CNS shows that the degree of its compactness is between octopus (compact CNS) and squid (elongated CNS) (see Fig 2.2 in Nixon and Young (2003)). Previous studies also suggested a high degree of similarity in CNS layout and underlying neural network between squid and cuttlefish, including the first of these from Cajal (1917), that initially highlighted the sophisticated visual and chromatophore systems (Sanders and Young, 1940, Boycott, 1953, Boycott, 1961, Messenger, 1968, Young, 1974, Young, 1976, Young, 1977, Young, 1979, Messenger, 1979, Budelmann and Young, 1987, Wild et al., 2015, Ponte et al., 2021). These works demonstrate the closeness of sepioids (cuttlefish) and teuthoids (squid) in spite of their long evolutionary separation (Strugnell et al., 2006, Allcock et al., 2014).

**Table 1.** List of ecological, behavioural, neuroanatomical features and estimates of lobe volume of decapodiforms used in this study. B- bean- shaped; C- croissant-shaped; D- diurnal; N- nocturnal.

| Species | Body size<br>Mantle length or<br>total length (mm)<br>(maturity) | OPs/CNS<br>(VL/CNS) | OP shape | Life mode | Habitat/depth | References |
| --- | --- | --- | --- | --- | --- | --- |
| *Sepia plangon* | ML 71-107<br>(adults) | 81% (4.3%) | C | D | Reef (down to<br>83m) | Current study |
| *Sepia latimanus* | ML 35<br>(40d post hatchling) | 83% (ca2%) | C | D | Reef (down to<br>30m) | Ziadi-Kunzli et al, 2018<br>Roper et al 2005 |
| *Sepia bandensis* | TL 80<br>(adult) | 74% (5.1%) | B | N | Reef | Montague et al 2022<br>Roper et al 2005 |
| *Sepia elegans* | - | 60% (3.2%) | B | - | Down to 500 m | Ziegler et al 2018<br>Roper et al 2005 |
| *Sepia officinalis* | ML 80<br>(subadult)<br>- | 67% (0.3%)<br>67% (3.7%) | B | N | Down to 200 m | Wirez 1959,<br>Denton & Gilpin-Brown, 1961<br>Maddock & Young, 1987<br>Roper et al 2005 |
| *Sepia omani* | TL 247<br>(adult) | -/- | B | - | 50-210m | Roper et al 2005 |
| *Sepia orbignyana* | - | 58% (3.3%) | B | - | 15-570m | Roper et al 2005 |
| *Sepia pharaonis* | ML 10 - 302<br>(hatchling to adult) | -/- | B | N | Down to 130m | Liu et al 2017<br>Roper et al 2005 |
| *Sepiella japonica* | - | -/- | B | - | 50m | Li et al 2018<br>Roper et al 2005 |
| *Sepioteuthis lessoniana* | ML 40 - 113<br>(juvenile) | 80% (2.6%) | B | cathemeral | Reef (down to<br>100m) | Chung et al 2020<br>Roper et al 2005 |

Early work on the organisation of the cuttlefish sensory and motor control systems was achieved through two methods: (1) Electrical stimulation of selected brain regions to detail the associating responses (Sanders and Young, 1940, Boycott, 1961, Chichery and Chanelet, 1976, Chichery and Chanelet, 1978, Chichery and Chichery, 1987). (2) Comparative studies in behavioural changes and learning impairment before and after brain region ablation (Sanders and Young, 1940, Boycott and Young, 1950, Chichery and Chichery, 1987).

The cuttlefish CNS was divided into 5 major functional regions: (i) The vertical lobe complex located at the most dorsal part of CNS with the noticeable dome-shaped vertical lobe (VL) and the superior frontal lobe (learning and memory). (ii) A pair of optic lobes (OPL) (visual tasks). (iii) A pair of peduncle lobes (PED) (cerebellum-like lobe for visual-motor control). (iv) Supra-oesophageal mass (Higher motor centres coordinating sensory inputs and behavioural responses). (v) Sub-oesophageal mass (Lower motor centres executing movement of funnel, arms, and mantle activities). This pioneering work produced a useful model for several ensuing studies of sensory reception, learning and memory (Messenger, 1973, Darmaillacq et al., 2006, Jozet-Alves et al., 2013, Yang and Chiao, 2016, Feord et al., 2020, Schnell et al., 2021b, Schnell et al., 2021a, Osorio et al., 2022).

Over the past two decades, a growing number of studies have focused on the behavioural neurobiology of the cuttlefish and their remarkably rapid and apparently cognitively complex reactions to novel challenges. For instance, cuttlefish can utilise spatial learning to solve maze tasks based on visual cues (e.g. landmark and e-vector of polarization light) (Alves et al., 2007, Cartron et al., 2012). Object recognition in cuttlefish (e.g. visual equivalence, amodal completion and visual interpolation for contour completion) appears to use strategies close to those used in vertebrates (Zylinski et al., 2012, Lin and Chiao, 2017a, Lin and Chiao, 2017b). The recent push towards comparisons of advanced cognitive behaviours (i.e. number sense, episodic-like memory, self-control), has postulated that the ability of the cuttlefish in solving complex tasks and cognitive reactions approaches that of young humans (Yang and Chiao, 2016, Schnell et al., 2021a, Schnell et al., 2021b).

Our current knowledge of the apparently complex behaviour of cuttlefish is predominantly derived from a large number of studies on a primarily nocturnal species, *S. officinalis* (Cajal, 1917, Sanders and Young, 1940, Boycott, 1961, Denton and Gilpin-Brown, 1961, Messenger, 1968, Chichery and Chichery, 1987, Nixon and Mangold, 1998, Gaston and Tublitz, 2004, King et al., 2005, Hanlon et al., 2009, Wild et al., 2015, Oliveira et al., 2017, Gonzalez-Bellido et al., 2018, Feord et al., 2020, Schnell et al., 2021a, Schnell et al., 2021b, Osorio et al., 2022). Despite intense interest their cognitive abilities the CNS gross anatomy, lobe organisation, brain-wide neural networks and the associated functional circuits is scant compared to both octopuses (Messenger, 1967, Young, 1971, Budelmann and Young, 1985, Plän, 1987, Chung et al., 2022) and loliginid squids (Cajal, 1917, Young, 1974, Young, 1976, Young, 1977, Young, 1979, Messenger, 1979, Wild et al., 2015, Chung et al., 2020).

Notably, while some of what we know around biology, ecology and physiology has also been obtained from the Indo-Pacific species, knowledge of their neuroanatomy is either sparse (e.g. *S. latimanus*, *S. pharaonis*, *Sepia bandensis* and *Sepiella japonica*) or absent among distinctively diurnal species such as *S. apama,* the flamboyant cuttlefish (*Metasepia pffeferi*) and the mourning cuttlefish (*Sepia plangon*) (Norman et al., 1999, Hanlon et al., 2007, Zylinski et al., 2011, Lee et al., 2013, Yang and Chiao, 2016, Liu et al., 2017a, Li et al., 2018, Schnell et al., 2019, Mezrai et al., 2020, Lu and Chung, 2017, Montague et al., 2022).

Recently developed techniques in magnetic resonance imaging (MRI) and histology to investigate cephalopod brains has revealed numerous novel findings at the morphological level. In particular, we have linked lobe growth and heterogeneity to ecological niches and lifestyle (Chung and Marshall, 2017, Liu et al., 2018, Chung et al., 2020, Chung et al., 2022).

Diffusion MRI (dMRI) using an ultra-conservative level for tractography acceptance has accurately delineated several new neural interconnections and networks, and at a level of detail not possible to see with conventional histology (Chung et al., 2020, Chung et al., 2022). It is worth noting that the first brain-wide connectome of squid CNS recovered 99.65% of the previously known neural tracts of loliginids (281 of 282) along with additional dozens of previously unknown visual-motor related tracts (Chung et al., 2020).

Furthermore, in contrast to a regular dorsoventral chiasmata in nocturnal octopuses, a new form of retinal wiring of the diurnal reef octopus which splits the visual scene into 4 separate zones suggested that this adaptation was linked to their ecology and behaviour (Chung et al., 2022). These examples highlight the advantage of new MRI-based methods and how a comparative study of various species, outside the list of the classical model species, allows evolutionary history to be drawn that may otherwise remain obscured.

In this context we asked three questions here: (1) Whether the neural anatomy of *S. officinalis* may be representative of all or most cuttlefish (over 100 species)? (2) Whether the cuttlefish brain may have some adaptations in response to their habits and habitats similar to those found in octopuses (i.e. enlargement and division of their visual centre, structural foldings and complexity in the learning and memory centre)? (3) Alternatively, given their free-swimming mode, are their brain adaptations more akin to their apparently closer cousins, the squid?

Understanding the gross neuroanatomy and circuit diagrams of any nervous system is the necessary first step towards understanding how evolution has shaped both brain structures and behaviours in cephalopods (Budelmann and Young, 1987, Nixon and Young, 2003, Williamson and Chrachri, 2004, Chung and Marshall, 2014, Chung and Marshall, 2017, Liu et al., 2018, Chung et al., 2020, Chung et al., 2022). In order to describe the neuroanatomy of the cuttlefish species described here, we have used the previous publications of *S. officinalis* and loliginid squids as a ‘baseline’, along with the few other descriptions for some brain areas that exist for other cuttlefish species (Cajal, 1917, Boycott, 1961, Young, 1974, Young, 1976, Young, 1977, Young, 1979, Messenger, 1979, Dubas et al., 1986b, Dubas et al., 1986a, Budelmann and Young, 1987, Wild et al., 2015, Liu et al., 2017a, Gonzalez-Bellido et al., 2018, Li et al., 2018, Chung et al., 2020, Montague et al., 2022).

We also chose a comparative approach, both between cuttlefish species and with squid, and investigated 2 species of decapodiform cephalopods that represent phylogenetically distinct groups and that exhibit different life modes, including the reef squid *Sepioteuthis lessoniana* and the diurnal reef cuttlefish, *S. plangon.* In addition to these species described here, another 9 cuttlefish species *(Metasepia tullbergi*, *Sepia elegans, Sepia orbignyana, Sepia omani*, *S. latimanus*, *S. officinalis, S. pharaonis, S. bandensis, S. japonica*) were selected from published literature (Boycott, 1961, Jereb and Roper, 2005, Wild et al., 2015, Liu et al., 2017a, Li et al., 2018, Ziegler et al., 2018, Montague et al., 2022) and included for further analyses where comparative data exists. Observations on the relative enlargement of brain lobes, and brain folding are included in an extended comparison of species, relative to ecology and lifestyle as well as phylogenies mostly based on existing morphological and molecular data.

## Results

### Gross anatomy of the diurnal cuttlefish brain

Dissection, contrast-enhanced 16.4T MR images (isotropic resolution 30 μm) and resulting 3D reconstruction show that the brain of *S. plangon* is located just under the anterior projection of the cuttlebone (Figure 1). The central complex (CC) is encased by the cranial cartilage whereas the two optic lobes (OPLs) are partially covered at the posterior end (Figure 1B). In gross anatomical terms this diurnal cuttlefish possesses a compact brain superficially similar to those of *S. officinalis* (histology and MRI (3T & 9.4T) (Tompsett, 1939, Boycott, 1961, Wild et al., 2015, Ziegler et al., 2018) and *S. bandensis* (histology and MRI (9.4T)) (Montague et al., 2022) and shares a similar lobe arrangement as the loliginid squids (Young, 1974, Young, 1976, Young, 1977, Young, 1979, Messenger, 1979, Chung et al., 2020), including 32 lobes (15 of which are bilateral) (Figures 1-2 & Table S1).

**Figure 1.**
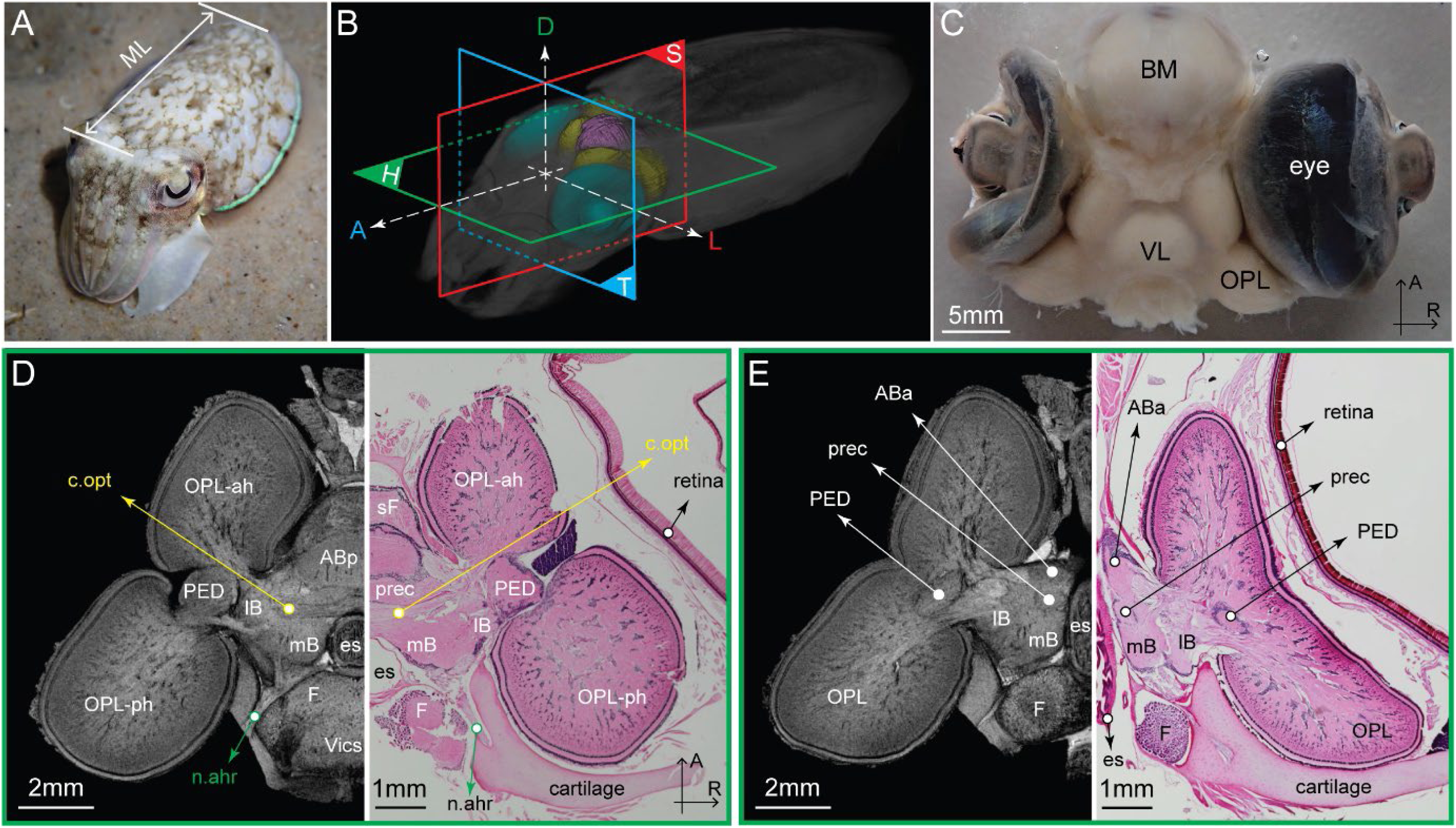
The diurnal cuttlefish, *Sepia plangon*, and the features of its central nervous system (CNS). (**A**) Live juvenile, *S. plango*n. ML - mantle length. (**B**) Three anatomical planes and 3D MRI rendering of an entire cuttlefish and the underlying CNS and eyes. H- horizontal; S- sagittal; T- transverse plane. A - anterior; P - posterior; D - dorsal; L – left; R - right lateral side. (**C**) Isolated brain-eyes preparation (dorsal view). BM- buccal mass; OPL - optic lobe; VL- vertical lobe. (**D-E**) Comparisons of horizontal sections between magnetic resonance histology (left) (isotropic resolution 30 μm) and conventional histology (right) (10 μm slice stained with hematoxylin and eosin). es- esophagus; Anterior anterior basal lobe (aBa); anterior posterior basal (aBp); optic connective (c.opt); anterior head retractor nerve (n.ahr); superior frontal (sF); lateral basal (lB); median basal (mB); precommisural (prec); peduncle (PED); fin (F); visceral (Vics); anterior horn of optic lobe (OPL-ah); posterior horn of optic lobe (OPL-ph).

**Figure 2.**
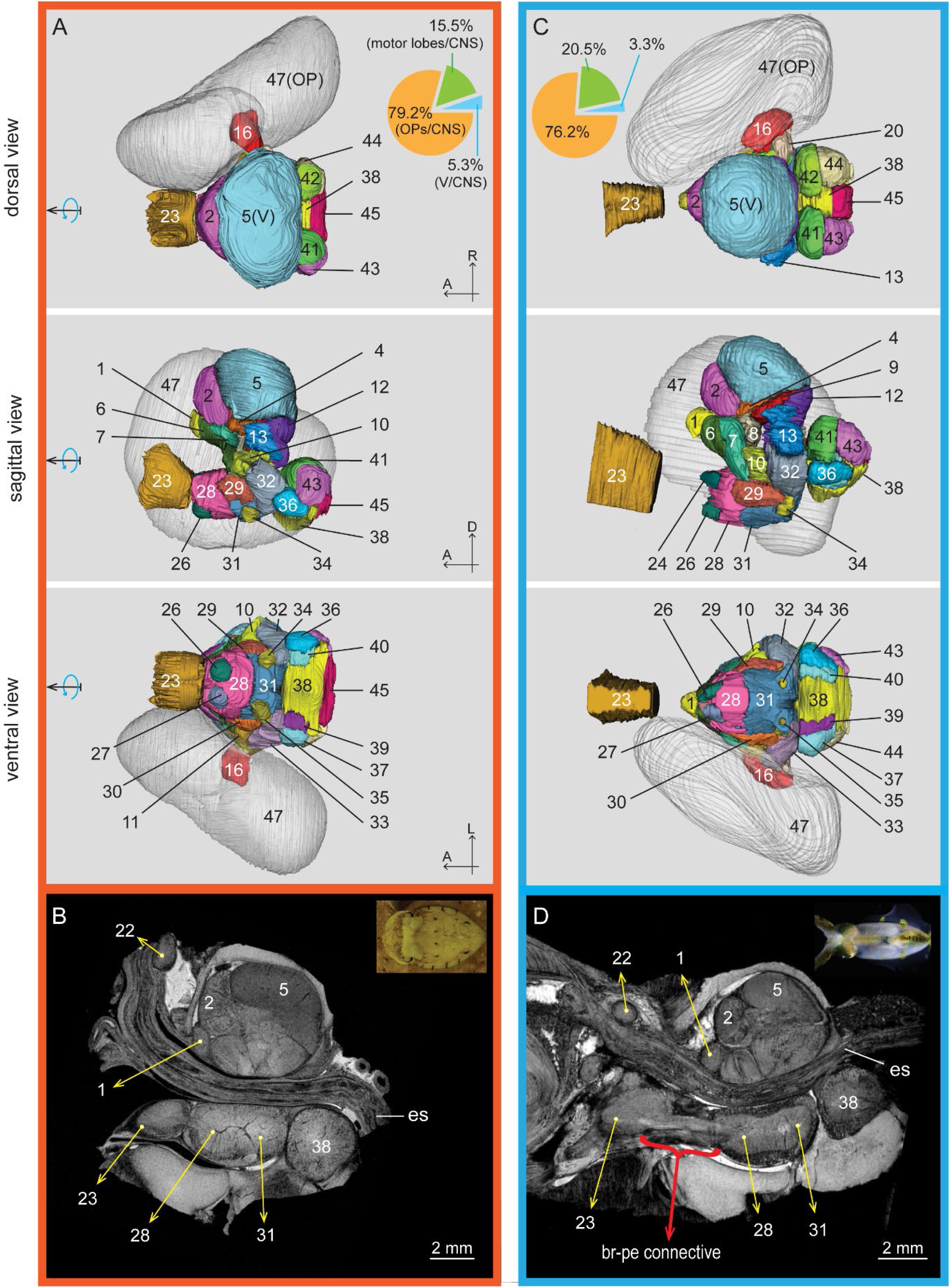
MRI-based 3D reconstruction of two types of decapodiform multi-lobed brains. (Top, middle and bottom rows are dorsal, sagittal, ventral viewpoints and sagittal section along the central midline). CNS gross anatomy and lobe organisation are superficially similar between cuttlefish and squid. (**A-B**) The diurnal tropical cuttlefish, *Sepia plangon*, its CNS layout and lobe-type are similar to that of the nocturnal temperate *Sepia officinalis*. (**C-D**) The reef squid *Sepiotetuhis lessoniana* and its CNS layout. (**D**) The long brachio-pedal connective makes the squid brachial lobe further away from the pedal lobe complex, rendering an elongated sub-esophageal mass compared to it of the cuttlefish (**B**). In total 47 lobes are identified (15 of which are bilateral) (See also Tables S1-2): (1) inferior frontal lobe; (2) superior frontal; (3) posterior frontal; (4) subvertical; (5) vertical; (6) anterior anterior basal; (7) anterior posterior basal; (8) precommissural; (9) dorsal basal (10-11) interior basal; (12) median basal; (13-14) lateral basal; (15-16) peduncle; (17-18) olfactory; (19-20) dorsolateral; (21) inferior buccal; (22) superior buccal; (23) brachial; (24-25) anterior dorsal chromatophore; (26-27) anterior ventral chromatophore; (28) anterior pedal; (29-30) lateral pedal; (31) posterior pedal; (32-33) dorsal magnocellular; (34-35) ventral magnocellular; (36-37) posterior magnocellular; (38) palliovisceral; (39-40) lateral ventral palliovisceral; (41-42) fin; (43-44) posterior chromatophore; (45) visceral; (46-47) optic.

Notably, the suboesophageal mass of squid is elongated due to the long brachio-pedal connective to make contact with the brachial lobe further away from the pedal lobe complex (Figure 2). Additionally, the close to bottom dweller, *S. plangon*, and the water column dweller, *S. lessoniana*, possess relatively small chemosensory regions (inferior frontal lobe complex (iFLx)), approximately 0.3-0.5% of CNS volume, indicating that chemoreception is less important than for the entirely benthic octopuses (4-6%) (Maddock and Young, 1987, Chung et al., 2020, Chung et al., 2022). Several previously unknown neuroanatomical features, obvious at a gross anatomical level, were identified in *S. plangon*, including distinct enlargement of the OPL and vertical lobe, and morphological folding of the OPL as described next (Figures 1-2, Videos S1-2).

### Croissant-shaped optic lobe

All specimens (1 hatchling, 2 juveniles and 3 adults) examined here possess distinct enlarged OPLs (the percentages of OPLs relative to total CNS volumes range between 77-82%) which are close to another diurnal cuttlefish *S. latimanus* (ca 82%) (Ziadi-Kunzli et al., 2019) and those of loliginid squids (e.g. 80% of CNS in *S. lessoniana*; *Sepioteuthis sepioidea* (79%) and *Loligo forbesi* (77%)) (Maddock and Young, 1987, Chung et al., 2020). This is in contrast to the moderately-large OPLs (58-74% of CNS) in another 4 cuttlefish species which are frequently active at low light conditions (Tables 1 & S3).

Another unique neuroanatomical feature of *S. plangon* among cuttlefish but one which it shares with some octopus species (Chung et al., 2022) is an only just described croissant-shaped OPL. All decapodiform cephalopods examined, as far as we know, have a regular bean-shaped OPL, including its cuttlefish siblings (e.g. *S. officinalis, S. bandensis, S. pharaonis, S. omani* and *S. japonica*), neritic squid (e.g. *Idiosepius*, *Loligo* and *Sepioteuthis*) and deep sea squid (e.g. *Abraliopsis*, *Architeuthis*, *Bathyteuthis*, *Liocranchia* and *Pyroteuthis*) (Boycott, 1961, Young, 1974, Chung, 2014, Chung and Marshall, 2017, Liu et al., 2017b, Liu et al., 2017a, Li et al., 2018, Liu et al., 2018, Montague et al., 2022).

Given their similar body size, the OPLs of *S. plangon* are significantly larger than those of the nocturnal *S. bandensis* (ML: 60-70mm) and the reef squid, *S. lessoniana* (ML: ca 110 mm) (Tables 1 & S2-3). The croissant-shaped OPL is present over a broad range of body size (young juvenile - adult, mantle length: 18-107 mm), less accentuated in the post-hatchling (a week old) and appears to be associated with a diurnal existence and associating visual tasks (Figures 2-4). Detailed morphological features are as follows:

(i) OPL horns. The dorsal 1/3 of the OPL is divided into two parts, forming two blunt horns that are closely opposed near the central line of the OPL. With the cuttlefish in a posture that is resting on the substrate or hovering in the water column, the anterior horn receives input from the posterior visual scene via the posterior vertical slit of its w-shaped pupil. The posterior horn is opposite to this and receives visual input from the anterio-ventral direction, a zone vital for the ballistic tentacular strike used for prey capture.
(ii) OPL sulcal folding. A second modification in *S. plangon* (again one found recently also in octopus (Chung et al., 2022)) is a curved-shaped sulcus at the lateral side apparently matched to the central crescent-shaped area of the pupil. The function of these structural folding is most likely to increase the surface area of the OPL. This is discussed relative to the gyrification index (GI =1.06) below but in brief appears to correlate with resolution power.

### Vertical lobe

Volumetric estimates show that the dome-shaped vertical lobe of *S. plangon* is significantly enlarged (4-5.3% of CNS volume) relative to those of the loliginid squids (0.3-3.2%) and cuttlefish species which are more active in the low light conditions (e.g. dominantly nocturnal *S. officinalis* (0.3-3.6%), and those living in deeper water (100-400m depth) such as *S. elegans* (3.2%) and *S. orbignyana* (3.3%)) (Table 1). Additionally, the size of vertical lobe increases significantly during ontogeny (from 2.4% at hatchling to approximately 4-5% at adult) (Tables 1 & S3).

### **T**ractography and connectome of the cuttlefish brain

Using the same imaging procedure and the selection criteria established for the squid brain (Chung et al., 2020), the averaged connectome of *S. plangon* (3 adults) allows recovery of all known major inter-lobed tracts described in squid and cuttlefish (n = 388, connectivity strength of tractography (*Cs* the logarithm of numbers of streamlines intersecting a pair of lobes: 0.48 - 5.76) (Figure 3) (Cajal, 1917, Boycott, 1961, Young, 1974, Young, 1976, Young, 1977, Young, 1979, Messenger, 1979, Budelmann and Young, 1987, Novicki et al., 1990, Chung et al., 2020). In addition, 181 blank spots (*Cs* = 0) in the averaged connectivity matrix from tractography are well-matched with the blanks from previous histology, demonstrating that our current procedure effectively eliminating false positives.

**Figure 3.**
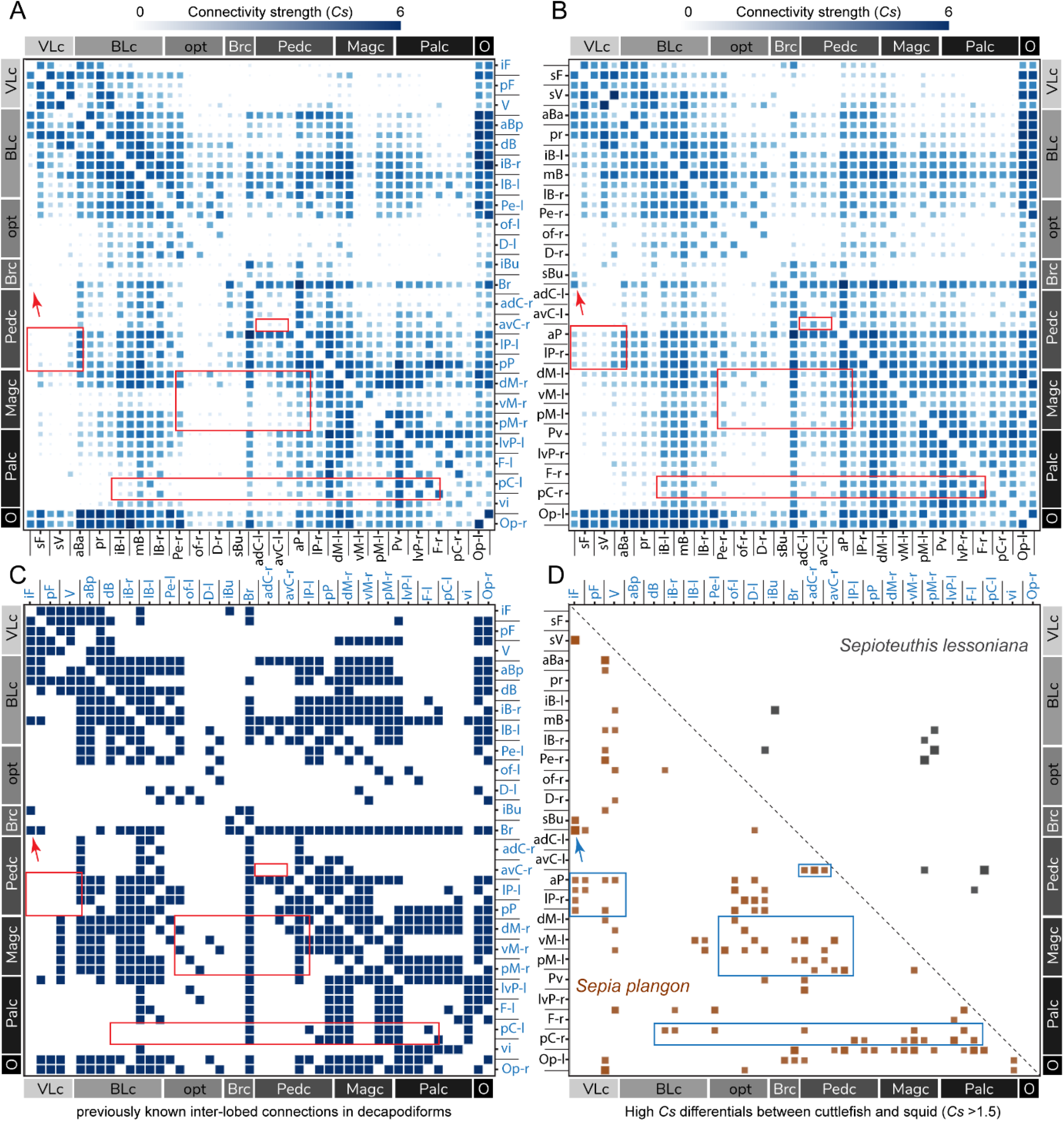
Comparisons of MRI-based connectivity matrices between the squid and cuttlefish brains. **(A)** An averaged probabilistic tractography connectivity matrix of *Sepioteuthis lessoniana* (5 juveniles). Arrow indicates distinct variation of the Cs value (brachial lobe - inferior frontal lobe) between two species. Another 4 highlighted regions showed significant variations between squid and cuttlefish. **(B)** An averaged probabilistic tractography connectivity matrix of *Sepia plangon* (3 adults). (**C**) This matrix summarized all described lobe-lobe neural connections of coastal decapodicforms (6 loliginid squid species and *Sepia officinalis*), including 388 neural connections (dark blue squares) based on a suite of silver impregnation and cobalt filling results initiated by J.Z. Young and his colleagues and our recent publication. **(D)** Using subtraction of the Cs values between two matrices, the major differences of connectivity strength (Δ*Cs* >1.5) between squid and cuttlefish were visualized. Brown squares represent that the inter-lobed connections are stronger in cuttlefish than those of squid. Grey squares are those strong connections in the squid brain lobes. The 4 highlighted regions show strong Cs values in cuttlefish which are related to chromatophore, magnocellular and pedal lobes where are potentially related to a large set of network in charge of complex colouration displays.

**Figure 4.**
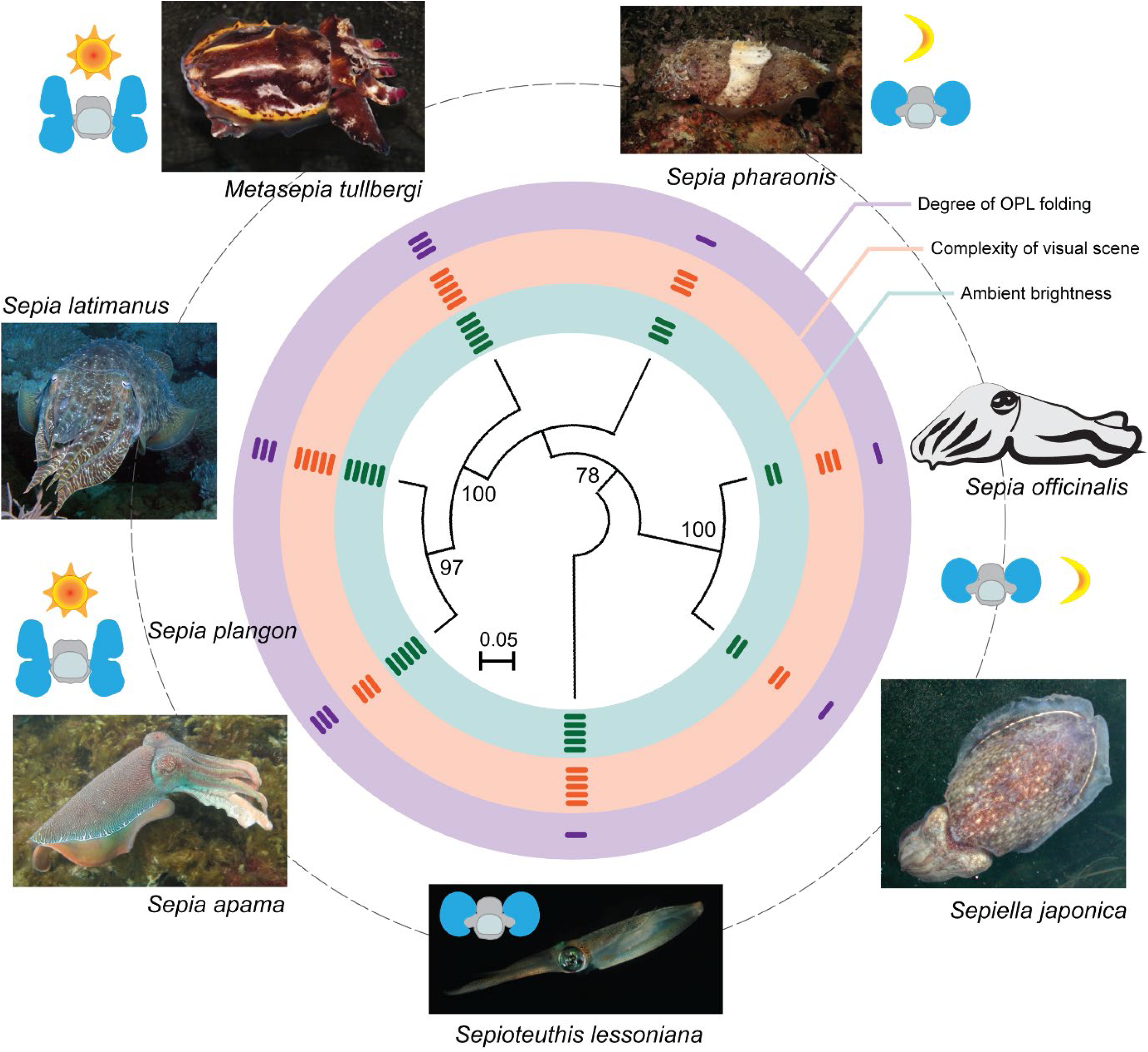
Diversity of neuroanatomical features in the optic lobe (blue) of the selected 7 decapodiforms and the corresponding life modes. The phylogenetical tree in the centre is constructed based on the published molecular information (entire mitochondrial DNA sequences were available in the selected 7 species, Star Method table). Due to partially sequenced molecular data, *Sepia plangon* was excluded. The bootstrap values are shown in front of the branch node. The neuroanatomical features and corresponding habit and habitats were based on the current study and the published literature (See also Table 1). Schematic sun indicates diurnal active species; schematic moons – nocturnal active species. Coloured bars in the two inner circles (green and orange) indicate degree of complexity of the visual scene estimated by their ecological niches and the ambient brightness the species inhabits based on published literature. Dark purple bars (in the purple circle) show the degree of structural folding of the optic lobe (OPL). A similar feature of the croissant-shaped OPL described in *Sepia plangon* is found in the three diurnal species using dissection.

Despite the considerable difference in phylogenetic relationship, a comparison of the MRI-based connectomes confirms a high degree of similarity in the inter-lobed network between squid and cuttlefish (Figure 3). Notably, the vision-related networks in two inter-lobed connectomes represent nearly the same pattern, including those connections between OPL-supra-esophageal mass (squid: 2.44 - 5.08 vs cuttlefish: 3.13 - 5.4) (e.g. OPL linked with basal lobe complex) and OPL-sub-esophageal mass with the median-high *Cs* value (squid: 0.74 - 4.58 vs cuttlefish: 1.18 - 5.21) (e.g. OPL linked with pedal and magnocellular lobes). Also, the connectomes within the sub-esophageal mass that are responsible for locomotion manoeuvre are similar, presumably due to similar modes of locomotion between the two groups. In addition, a comparison of the *Cs* between brachial and inferior frontal lobes (*S. plangon* (3.77) vs *S. lessoniana* (0.61)) confirms a previous qualitative description that a strong inter-lobed connection throughout the cerebro-brachial tracts exists in cuttlefish, *S. officinalis*, whereas fewer stained neurons are seen in squid, *Loligo vulgaris* (Budelmann and Young, 1987) (Figure 3).

A few remarkably strong inter-lobed connections (*Cs*: 1.6-2.8) may be identified as tracks unique to cuttlefish, whereas those in the squid connectome are either absent or with a much lower *Cs* value (<1), including those related to the chromatophore, magnocellular and pedal lobes (Figure 3D). Considering the main function of these three brain regions, in control of locomotion and colouration (Boycott, 1961), these previously-unknown tractographic connections are likely to drive dynamic body pattern changes as well as the two previously known circuits (OPL-lB-Ch and OPL-PED-lB-Ch) (Messenger, 2001, Gonzalez-Bellido et al., 2018).

### Phylogenetic analyses

Pagel’s λ and phylogenetic generalised least squares (PGLS) analyses were used to estimate the likelihood that these newly described modifications are phylogenetically linked (STAR Methods). A strong phylogenetic relationship is linked in the morphological changes of the optic lobes (Pagel’s λ= 0.9999 for all 7 species; test of λ = 1, p = 1) (Figure 4). However, we suggest the adaptations seen here, especially those within the OPL in diurnal reef dwellers are most likely driven by the needs of their life modes. In other animals, it is the adaptations of the central brains and existing CNS design that are more likely to retain a phylogenetically-flavoured relationship (Yopak et al., 2010, Yopak et al., 2020, Chung et al., 2022, Wolff et al., 2017, Nixon and Young, 2003).

## Discussion

In common with their major competitors, the fish, coastal cephalopods are successful and voracious visual predators that live over a broad range of ecological niches. In contrast to our knowledge of fish neuroanatomical adaptations related to sensory perception, foraging modes and habitats (Wagner, 2001, Lisney and Collin, 2006, Yopak et al., 2015), establishing links between behavioural features and neuroanatomical modifications remains in its infancy for the cephalopods (Ponte et al., 2021). Using MRI-based techniques and conventional histology, we have started the first detailed comparison of neuroanatomical features and corresponding MRI-based connectomes between cuttlefish, octopus and squid. This work focusses on cuttlefish but uses our previous studies on squid and octopus as a comparison, both to describe differences in basic brain and wiring anatomy and to examine the ecology and, to an extent, the evolution of the cephalopod brain. It of course stands on the shoulders of previous work on these brainy invertebrates, notably that of JZ Young and colleagues (Boycott, 1961, Messenger, 1973, Nixon and Young, 2003) along with a few studies between that time and now (Chichery and Chanelet, 1976, Chichery and Chanelet, 1978, Chichery and Chichery, 1987, Dickel et al., 1997, Williamson and Chrachri, 2004, Liu et al., 2017a, Gonzalez-Bellido et al., 2018).

We also present new findings from a comparative approach amongst cuttlefish species and hope to provide a firm base to challenge the long-standing assumption that neuroanatomical features of *S. officinalis* are representative of all cuttlefish species. The neuroanatomical variation we note here infers interspecific variation in visual capabilities, the importance of vision relative to the less utilised chemosensory system and clear links with life modes (e.g. diurnal vs nocturnal), ecological factors (e.g. living depth and ambient light condition) as well as to an extent, phylogeny.

### Unique neuroanatomical features in the mourning cuttlefish, *Sepia plangon*

Early reports divided the cuttlefish brain into regions and associated functions based on electrical stimulation of selected lobes of *S. officinalis* (Boycott, 1961, Chichery and Chanelet, 1976, Chichery and Chanelet, 1978, Chichery and Chichery, 1987). However the neuronal number and circuitry behind these connections has remained largely unknown for now more than 30 years (Budelmann and Young, 1987). Here MRI-based observations and gross anatomy have revealed a number of new observations.

The tropical diurnal reef cuttlefish, *S. plangon*, apparently possesses an enlarged brain compared to the other coastal species with a similar given body size. The adult-like hatchling of *S. plangon* (ML: 8 mm) has an enlarged brain compared to *S. officinalis* (ML: 6.3 mm) (CNS: 9.26 vs 2.94 mm^3^ and OPLs: 7.11 vs 1.97 mm^3^) (Wild et al., 2015). Notably, the cuttlefish embryo starts to react to visual and chemical cues before hatching (stage 30) (Darmaillacq et al., 2006, Mezrai et al., 2020). Unlike the eggs of *S. officinalis* which are darkened by maternal ink resulting in poor visibility of the outside scene, the transparent egg of *S. plangon* allows the embryo to receive surrounding visual cues and respond accordingly with flashing chromatophores. This early vision-related demand toward the post-hatching environment may therefore initiate enlargement of the OPL of *S. plangon* more than that seen in *S. officinalis* (77% vs 67% of CNS).

The CNS of *S. plangon* grows rapidly and particularly the VL and OPLs attain a level of complexity and volume not seen in previously examined cuttlefish (Table S3). The size increase of VL during ontogeny results in a 210% relative increase from 2.4% of CNS at hatchling to approximately 4-5% at adult. Furthermore, growth of the OPL from post-hatchling to adult is up to 100 fold the volume increase during all life stages, emphasising the vital role of vision for this diurnal species.

Our examination of *S. plangon* shows, like octopus (Chung et al., 2022) two types of OPLs exist, bean vs croissant shape and that this reflects their phylogenetic relationship, life modes and habitats (Figure 4). Both nocturnal and deep-water dwelling cuttlefish species (>200m depth) which encounter dim light condition have a regular bean-shaped OPL (Table 1). In contrast, the diurnal species seem to have the enlarged croissant-shaped OPL, a modification associated with a more visual existence and first noted in our previous studies on diurnal octopus species octopus (Chung et al., 2022). By contrast, cuttlefish that live in low light condition where there is less visual contrast possess smaller OPL than those of the diurnal species (Ziegler et al., 2018) (Figure 4).

### Similarity of brain regions between squid and cuttlefish

Squid and cuttlefish predation is remarkably fast and precise. The feeding behaviour entails a rapid tentacular strike to catch small prey and a ‘punch’ from the arm crown to attack and defend for large objects (Chung and Marshall, 2014, Lu and Chung, 2017, Hanlon and Messenger, 2018). These ballistic movements are visually-coordinated activities and include finding a prey item in the distance and, on moving closer, estimating the object size to guide ballistic strike (Messenger, 1968, Kier and Von Leeuwen, 1997, Chung and Marshall, 2014, Feord et al., 2020). Additionally, assessment of prey quality (acceptation or rejection for feeding) is based on contact chemoreception via the suckers of the arms and tentacles (Messenger, 1973, Archdale and Anraku, 2005).

The proportion of neural processing investment in chemoreception and vision between the three coleoid groups (cuttlefish, squid and octopus) is quite variable and this study has helped uncover new and underline previous observations. All three cephalopod groups possess optically excellent and often large eyes and all three put considerable investment into the OPL processing of vision (but see ecological differences discussed in Chung et al. (2022)) (Land, 1981, Sweeney et al., 2007). There is a difference in volume ratio between the two sensory brain regions, vision (OPLs) versus chemoreception (iFLx), which reaches over 100 fold in cuttlefish (e.g. 101 in *S. officinalis*; 235 in *S. plangon*), > 200 in loliginid squid (e.g. 220 in *S. lessoniana*; 305 in *Loligo forbesi*) compared to a very low value around 10 in the benthic octopuses, such as *Octopus vulgaris* and *Hapalochlaena fasciata* (Maddock and Young, 1987, Chung et al., 2020, Chung et al., 2022). The relative value of a given sensory area clearly shows its level of importance, suggesting again that the water column dwellers rely more on vision, whereas the more benthic groups favour a combination of vision and chemoreception.

Further to the basic volumetric data, vision-related connectomes highlight that cuttlefish and squid have adopted similar principles in design in response to visually-coordinated activities at a very fine scale (Figures 2-3). These two groups possess similar network matrices within the vision to higher motor brain regions (e.g. basal lobe complex) (Figure 3).

Again vision related, the multilayered structure in all basal lobes show tractographic projections from the upper layers of the basal lobes that connect only with the upper level of the optic lobe, whereas the projections from the lower levels of the basal lobes shift toward lower levels of the optic lobe accordingly (Chung et al., 2020) (Video S3). This multi-layered network arrangement likely retains retinotopic spatial information from the outside world through to the motor command units in the BLs (Young, 1977, Chung et al., 2020).

Finally, this direct connection from visual input in to motor action out is underlined by the new finding that the basal lobe complex possesses interweaving circuits with the sub-esophageal mass. This suggests a relay station exists, mediating motor control such as arm movements (brachial lobe), tentacular strike and eye movements (pedal lobes) and funnel and fin movements (magnocellular, fin and palliovisceral lobes) (Boycott, 1961, Young, 1976, Chichery and Chichery, 1987, Budelmann and Young, 1987, Chung et al., 2020).

### Cuttlefish-unique neural network features related to chemoreception, colouration and camouflage?

While there is a degree of similarity in inter-lobed connectivity between cuttlefish and squid brains, there are also other tractographic, network and gross anatomical features unique to cuttlefish. These again appear largely driven by ecology and behavioural habits. In brief they are the network between iFLx and brachial lobe (chemosensory related circuits) and those amongst chromatophore, magnocellular and pedal lobes (colouration related circuits) (Figure 3). Each of these cuttlefish-unique features is now described in more detail based around suggested function.

### Chemosensory-learning circuits

At gross anatomic levels, the volumetric ratio between iFLx and OPLs in squids is smaller than in cuttlefishes in both temperate (e.g. *L. vulgaris* vs *S. officinalis*) and tropical (e.g. *S. lessoniana* vs *S. plangon*) species (Maddock and Young, 1987, Chung et al., 2020). In addition, the increasing complexity of neural network between brachial lobe and iFLx in cuttlefish indicate that cuttlefish may favour chemosensory cues in daily tasks and more so than squid (Figures 2-3). In the behavioural context, bait coated with additional chemicals or biological extract (e.g. amino acids, quinine or cephalopod ink), may be accepted or rejected by touching the bait using arms/tentacles in the cuttlefish, *Sepia esculenta* (Archdale and Anraku, 2005). A similar bait handling behaviour has been found in the other 2 cuttlefish, *S. plangon* and *S. latimanus*, during feeding training in captivity. Using the same method rarely triggered feeding acceptance by squid such as *S. lessoniana* that appear to need movement cues to trigger bait capture (personal observation). This indicates that cuttlefish maintains good contact chemosensory capabilities, somewhere between octopus and squid, which could be helpful to shape prey preference and tune foraging strategies.

### Additional colouration related circuits in cuttlefish

Numerous novel projectomes related the cuttlefish chromatophore lobe (Cs > 1.5) are identified in the matrix (Figure 3D). Although the function of this network remains unclear, two possible explanations are proposed as follows: (1) Ontogenetic differences. (2) Additional circuits related to body patterns.

#### (1) Ontogenetic differences

The cephalopod brain grows continuously over a long period time during its limited 1-2 year life span. This is accompanied by an increasing complexity of behaviours (Messenger, 1973, Nixon and Young, 2003, Chung et al., 2020, Chung et al., 2022). For instance, the hatchling of *S. plangon* only shows two simple body patterns (uniform darking and blanching) in contrast to the diverse colouration displays during courtship and sophisticated camouflage and warning patterns (Alejandra et al., 2020). Considering the current squid connectome based on 5 juveniles (ML: 40-113 mm) along with other supporting neural tracing data that also favoured smaller brains (mainly juveniles) (Young, 1976, Budelmann and Young, 1987, Novicki et al., 1990, Chung et al., 2020), a comparison between the two connectomes (adult cuttlefish vs juvenile squid) could therefore miss some connections which appear only at the adult stage.

#### (2) Additional circuits related to cuttlefish body patterns

Decapodiform cephalopods show several forms of courtship display which visually attract mates and coordinates copulation activities (Brown et al., 2012, Lin et al., 2017, Hanlon and Messenger, 2018, Alejandra et al., 2020). Cuttlefish courtship display has been well documented in a few species, including *S. latimanus*, *S. officinalis* and *S. plangon.* These displays often use a combination of chromatic, textural and postural components (Hanlon and Messenger, 2018, Alejandra et al., 2020). For instance, *S. plangon* uses 34 chromatic components combined with 3 textural and 14 postural components for dynamic courtship displays (11 patterns used by female; 18 by male) (Alejandra et al., 2020). In contrast, squid mainly relies on chromatic components alone such as *S. lessoniana* assembling 27 chromatic components during reproductive interactions (7 patterns used by female; 12 by male) (Lin et al., 2017). It should be remembered that both groups are most likely colour blind, seeing only the luminance and pattern component of such displays (Marshall and Messenger, 1996, Chung and Marshall, 2016).

The complexity of camouflage tricks between cuttlefish and squid is also substantial. Cuttlefish camouflage contains a combination of cryptic colouration, skin texture and arm posture to conceal itself into the 3D characters of the surrounding scene (e.g. algae, rubbles, coral) (How et al., 2017, Gonzalez-Bellido et al., 2018, Hanlon and Messenger, 2018). By contrast, the squid mainly relies on colour changes on body surface to mimic the 2D background such as manipulating colours to match with substrate while reaching close to floor and switching to countershading while hovering in water column (e.g. *S. lessoniana*) (Lu and Chung, 2017, How et al., 2017, Nakajima et al., 2022). Both chromatic and hydrostat systems are regularly used in the formation of cuttlefish body patterns (Gonzalez-Bellido et al., 2018, Alejandra et al., 2020, Osorio et al., 2022), and one additional set of neural components to coordinate those apparently more complex body patterns compared to a relatively simple system used for the squid chromatic-based patterns is revealed here (Figure 3). The detailed function will need further tests to clarify what these additional circuits achieve relative to neural and behavioural dynamics and how the cuttlefish nervous system dispatches signals via different pathways to govern skin patterns (Laan et al., 2014, Reiter et al., 2018, Osorio et al., 2022).

### Elongated CNS layout linked to the streamline body shape

3D reconstruction of the coastal decapodiform brain clearly showed that distinct CNS elongation appears in the myopsid squid and not in cuttlefish (Figure 2). Firstly, with the absence of a floatation apparatus, the cuttlebone, to offset gravity, squid rely on constant swimming to maintain buoyancy and direction, resulting in a daily energy cost much higher than that of the neutral buoyant cuttlefish (O’Dor, 2002). This means that a long, streamlined body shape that minimises energy consumption is desirable for squid (O’Dor and Webber, 1986). In turn this has resulted in a stretched squid brain, to fit within this body shape and prevent its brachial and optic lobes bulging outward, causing higher drag. A similar observation of a further elongated CNS layout was briefly described in the oceanic oegopsid squid (neon flying squid) by Nixon and Young (2003)), again suggesting that development of the streamline body shape of squid might be therefore co-evolved with its elongated CNS.

## Materials and Methods

### Sample collection and preparation

All collections were conducted under a Great Barrier Reef Marine Park Permit (G17/38160.1) and Queensland General Fisheries Permit (180731). The mourning cuttlefish, *Sepia plangon*, and oval squid, *Sepioteuthis lessoniana*, were collected using a seine net (water depth 1-3m) close to Moreton Bay Research Station, Stradbroke Island, Queensland, Australia. The maintenance and experimental protocol used here were covered by animal ethics permit (QBI/236/13/ARC/US AIRFORCE & QBI/304/16). Total 44 cuttlefish and 5 squid were collected for this neuroanatomical study in 2017-2021.

Animals were anaesthetised in cool seawater (15°C) mixed with 2% MgCl_2_ (Chem-Supply, Australia) and sacrificed by an overdose of MgCl_2_ prior to fixation. The small specimens (hatchlings and early juvenile) were soaked into 4% PFA-PBS fixative at 4° C for 48 h and then transferred to 0.1% PFA-PBS fixative for storage at 4 °C until further dissection.

Three adult cuttlefish specimens for MR imaging were fixed using the transcardial perfusion protocol developed by Chung et al. (2020). In brief, the transcardial perfusion protocol is using 4% paraformaldehyde (PFA) (EM grade, Electron Microscopy Sciences, Hatfield, USA) mixed with 0.1 M PBS with the rate of perfusion set to 2.5 ml per minute. The perfusion proceeded until 0.2 ml fixative per gram of specimen was used. Subsequently the muscle, skin and connective tissues around the brain were removed and the specimen was soaked in 4% PFA-PBS fixative for overnight to reduce morphological deformation of the brain.

### Image stacking of the isolated brain-eyes

The isolated brain and eyes were imaged with the focus stacking method using a digital camera (Canon 5D4 camera with Canon MPE 65mm Macro lens, Canon, Japan) mounted on the electronically-controlled focusing rack (Castel-Micro focusing rack, Novoflex, Germany). A sequence of close-up images was captured from the dorsal end of brain to the ventral end using 0.1 mm step for small samples or 0.25 mm step for large samples. Focus stacking (20-80 images) was processed using the software Helicon Focus Pro (version 7.6.4, Helicon Soft Ltd. Ukraine), rendering an image with a greater depth of field.

### MRI procedure

Intact brain and eyeballs were isolated and repeatedly rinsed with 0.1 M PBS to minimise fixative residue. The isolated brain and eyes were then soaked into 0.1 M PBS containing magnetic resonance imaging (MRI) contrast agent, 0.2% ionic Gd-DTPA (Magnevist) (Bayer, Leverkusen, Germany), for 24-48 hours to enhance image contrast prior to MR imaging (Chung et al., 2020, Chung et al., 2022). Six contrast-enhanced cuttlefish brains were imaged following the protocol developed by Chung et al. (2020). The contrast-enhanced specimen was placed into a fomblin-filled (Fomblin oil, Y06/6 grade, Solvay, USA) container to prevent dehydration and then vacuumed for 3 minutes to remove air bubbles trapped inside oesophagus or brain lobes. The container was then placed in a custom-built 20 mm diameter surface acoustic wave coil or 10 mm diameter quadrature coil (M2M Imaging, Brisbane, Australia). Both high resolution MR structural images and high angular resolution diffusion images (HARDI) were acquired using a 16.4 Tesla (700 MHz) vertical wide-bore microimaging system (interfaced to an AVANCE I spectrometer running imaging software Paravision 6.0.1 (Bruker Biospin, Karlsruhe, Germany) in the Centre for Advanced Imaging at the University of Queensland. Imaging was performed at a room temperature (22 °C) using a circulating water-cooling system.

Three dimensional (3D) high resolution structural images were acquired using fast low angle shot (FLASH) with the following parameters based on Chung and Marshall (2017): echo time (TE) / repetition time (TR) = 12/40 ms, average = 4, flip angle (FA) = 30°, field of view (FOV) = 7.5 × 6.4 × 6 mm to 21 × 13 × 13 mm for different individuals, 30 μm isotropic resolution. Total acquisition time for one brain was 1 h (hatchling) to 8.3 h (the largest brain).

After FLASH imaging, 3D high angular resolution diffusion-weighted imaging (HARDI) was acquired with the following parameters: TR = 300 ms, TE = 22 ms, 30 direction diffusion encoding with b-value = 3000 s/mm^2^, two b0 images acquired without diffusion weighting and 80 μm isotropic resolution with 1.5 partial Fourier acceleration acquisition in the phase dimensions (Chung et al., 2020). Total acquisition time for one brain was 16.5-35.5 h.

### Estimates of lobe volume

Identification of the cuttlefish brain lobes was based on the published anatomical studies of cuttlefish and loliginid squids as an initial aid in determining the boundaries between tissue. 47 lobes previously defined by (Young, 1974, Young, 1976, Young, 1977, Young, 1979, Messenger, 1979, Chung et al., 2020, Boycott, 1961) were identified from the MRI data. The parcellation of the selected lobes and brains was then manually segmented using MRtrix3 (version 3.0.2, open-source software, http://www.mrtrix.org/) (Tournier et al., 2019) and then estimates of volume of the selected lobes and an entire brain were calculated using ITK-SNAP (version 3.6.0, open-source software, http://www.itksnap.org/) (Yushkevich et al., 2006). Considering variations of volume estimates of cephalopod brain which are strongly affected by the size and age of the individuals, the volumes of the lobes were expressed as percentages of the total CNS volume to circumvent this issue as suggested in previous studies (Maddock and Young, 1987, Chung et al., 2020, Chung et al., 2022).

### Construction of structural neural connectivity matrix

Our previous work demonstrated that the high resolution HARDI combined with conservative selection criteria enabled to accurately reveal the major neural tracks in the squid brain and octopus optic nerve tracks (Chung et al., 2020, Chung et al., 2022). Adapting the same procedure to construct the brain-wide tractography of cuttlefish brain, the 47 lobes, regions of interest (ROIs) were used to construct tractography. Probabilistic fibre tracking was then performed using second order integration over the fibre orientation distribution (FOD) algorithm and the tracts were generated independently for each ROI (10 streamlines per voxel) with an optimized FOD amplitude cut-off value of 0.175 to generate biologically realistic tractography in cephalopod neural tissue at mesoscale. The brain-wide cuttlefish neural connectivity matrix where the connections and the corresponding connectivity strength (*Cs*) were mapped to the relevant cuttlefish brain lobes for each individual. The averaged pairwise *Cs* were also calculated and plotted in the matrices for further analysis with the previously-published squid matrix (Chung et al., 2020).

### Contour-based measurement of gyrification index (GI)

The degree of folding of the optic lobe was measured using the contour-based method (Chung et al., 2022). We measured the GI by comparing the lengths of complete and outer contours of the selected brain lobes in a serial horizontal MR slices for the OPLs along with the dorso-ventral axis using Fiji (version 1.53c, open-source software, https://imagej.net/) (Schindelin et al., 2012). The mean GI of the defined entire lobe is the ratio between the sum of the total outer contour and the sum of the superficially exposed surface contours.

### Phylogenetic analyses

In order to understand whether the phylogenetic relationship or the life mode affect the modification of octopodiform’s brain, the phylogenetic generalised least squires (PGLS) test was used to investigate the impact of several predictor variables (life modes, light conditions, and visual tasks) on the modification of neuroanatomical structure while controlling for potential phylogenetic signals in the responses (Mundry, 2014). Determination of the selected octopus phylogenetic relationships was based on the published complete mitochondrial DNA sequence which were available from GenBank. Alignments of sequence were constructed using the multiple sequence alignment (MUSCLE) method with MEGA X (molecular evolutionary genetics analysis program version 10.2.5) (Kumar et al., 2018). *Sepioteuthis lessoniana* was used as the outgroup. The phylogenetic tree of these selected species was generated by the Maximum-Likelihood method and the bootstrap confidence values (1000 replicates) were calculated with MEGA X (Kumar et al., 2018). The phylogenetic signal was estimated with Pagel’s λ using the package the CAPER v1.0.1 as implemented in the RStudio v1.4.1103. The relationship between the changes of brain anatomy and environmental characters (Table S3) was determined using the phylogenetic generalised least squares (PGLS) method with the CAPER package in RStudio.

## Acknowledgements

This work is supported by the Australian Research Council (ARC) (Australian Laureate Fellowship (FL140100197) to N.J.M.), (Discovery Project (DP200101930) to N.J.M.) and the Office of Naval Research Global (ONR Global) (N62909-18-1-2134 to N.J.M.) The 16.4T is supported by the Queensland State Government through the Queensland NMR Network, and the Australian Government through National Collaborative Research Infrastructure Strategy (NCRIS) and the National Imaging Facility. We thank the staff of the Moreton Bay Research Station for logistical support. We also acknowledge the Quandamooka people as the Traditional Owners and their custodianship of the lands on which Moreton Bay Research Station operate. We pay our respects to their ancestors and their descendants, who continue cultural and spiritual connections to Country and recognise their valuable contributions to Australian and global society.

## Author contributions

Conceptualization, W.-S.C. and A.L.G.; methodology, A.L.G. N.D.K. and W.-S.C.; funding acquisition and supervision, N.J.M.; validation and visualization, W.-S.C. N.D.K. and N.J.M.; the first draft of manuscript, W.-S.C.; all authors contributed to data analysis, interpretation and revision of the manuscript.

---

**Table S1.** List of cuttlefish brain lobes with abbreviations used through the text The main functions of the lobe systems based on work by Young and his colleagues (Messenger, 1979; Young, 1961, 1971; 1974, 1976; 1977, 1979; Boycott and Young, 1955, 1957; Boycott, 1961; Nixon and Young, 2003). Supraoesophageal mass includes basal lobe and optic track complexes. Suboesophageal mass consists of the brachial lobe, pedal lobe, magnocellular lobe, and palliovisceral lobe complexes. * indicates that the lobe is further divided into the left and right lobe. ^Ψ^ indicates a further sub-division of the anterior chromatophore lobes into dorsal and ventral halves.

| Lobe system and function | Lobe | Abbreviation |
| --- | --- | --- |
| Vertical lobe complex (VLc)<br>- Memory & learning | Inferior frontal | iF |
|  | Superior frontal | sF |
|  | Posterior frontal | pF |
|  | Subvertical | sV |
|  | Vertical | V |
| Basal lobe complex (BLc) -<br>Higher motor control | Anterior anterior basal | aBa |
|  | Anterior posterior basal | aBp |
|  | Precommisural | pr |
|  | Dorsal basal* | dB |
|  | Interbasal* | iB |
|  | Median basal | mB |
|  | Lateral basal* | lB |
| Optic track complex (opt) -<br>Intermediate visual-motor center &<br>olfaction | Peduncle* | Pe |
|  | Olfactory* | of |
|  | Dorsolateral* | D |
| Brachial lobe complex (Brc) -<br>Arm and feeding control | Inferior buccal | iBu |
|  | Superior buccal | sBu |
|  | Brachial | Br |
| Pedal lobe complex (Pedc) -<br>Intermediate and lower motor<br>center for locomotion control | Anterior dorsal chromatophore* <sup>Ψ</sup> | adC |
|  | Anterior ventral chromatophore* <sup>Ψ</sup> | avC |
|  | Anterior pedal | aP |
|  | Lateral pedal* | lP |
|  | Posterior pedal | pP |
| Magnocellular lobe complex<br>(Magc) - Intermediate motor<br>center | Dorsal magnocellular* | dM |
|  | Ventral magnocellular* | vM |
|  | Posterior magnocellular* | pM |
| Palliovisceral lobe complex<br>(Palc) - Lower motor center for<br>locomotion and mantle activities | Palliovisceral | Pv |
|  | Lateral ventral palliovisceral* | lvP |
|  | Fin* | F |
|  | Posterior chromatophore* | pC |
|  | Visceral | vi |
| Optic lobes (O) - Vision | Optic* | OPL |

**Table S2.** Estimates of lobe volume of the mourning cuttlefish, S*epia plangon*

|  |  |  | Hatchling<br>ML:8mm | Juvenile<br>ML:18mm | Juvenile<br>ML:32.4mm | Adult male<br>ML:72.9mm | Adult female<br>ML:71.1mm | Adult female<br>ML:107mm |
| --- | --- | --- | --- | --- | --- | --- | --- | --- |
| VL complex | iFL | 1 |  |  | 0.11 | 0.36 | 0.39 | 0.97 |
|  | sFL | 2 |  |  | 1.14 | 3.56 | 3.67 | 8.18 |
|  | pFL | 3 |  |  | 0.16 | 0.13 | 0.20 | 0.16 |
|  | sVL | 4 |  |  | 0.95 | 3.43 | 4.26 | 9.04 |
|  | VL | 5 |  |  | 4.14 | 15.62 | 18.12 | 46.61 |
| ABL complex | ABL-a | 6 |  |  | 0.47 | 1.62 | 1.91 | 4.49 |
|  | ABL-p | 7 |  |  | 0.55 | 1.74 | 1.69 | 3.56 |
|  | precL | 8 |  |  | 1.02 | 1.48 | 2.41 | 3.14 |
| BL complex | dBL | 9 |  |  | 0.52 | 0.62 | 1.42 | 2.67 |
|  | intBL-L | 10 |  |  | 0.19 | 0.54 | 0.81 | 1.85 |
|  | intBL-R | 11 |  |  | 0.16 | 0.53 | 0.73 | 1.93 |
|  | mBL | 12 |  |  | 1.62 | 4.73 | 5.07 | 14.37 |
|  | IBL-L | 13 |  |  | 0.21 | 1.01 | 0.67 | 1.21 |
|  | IBL-R | 14 |  |  | 0.21 | 1.01 | 0.63 | 1.45 |
| OPT complex | PeduncleL | 15 |  |  | 0.31 | 1.00 | 1.32 | 2.54 |
|  | PeduncleR | 16 |  |  | 0.36 | 0.99 | 1.20 | 2.54 |
|  | ofL-L | 17 |  |  | 0.06 | 0.17 | 0.29 | 0.05 |
|  | ofL-R | 18 |  |  | 0.05 | 0.17 | 0.31 | 0.03 |
|  | DorsolatL-L | 19 |  |  | 0.05 | 0.16 | 0.29 | 0.18 |
|  | DorsolatL-R | 20 |  |  | 0.05 | 0.16 | 0.31 | 0.19 |
| BrachL complex | iBuL | 21 |  |  | 0.14 | 0.39 | 0.91 | 1.76 |
|  | sBuL | 22 |  |  | 0.22 | 1.05 | 1.11 | 2.00 |
|  | BrachiL | 23 |  |  | 0.87 | 5.45 | 6.40 | 12.21 |
| PedL complex | Chrom-dA-L | 24 |  |  | 0.08 | 0.31 | 0.32 | 0.28 |
|  | Chrom-dA-R | 25 |  |  | 0.08 | 0.32 | 0.31 | 0.29 |
|  | Chrom-vA-L | 26 |  |  | 0.09 | 0.32 | 0.26 | 0.48 |
|  | Chrom-vA-R | 27 |  |  | 0.08 | 0.30 | 0.29 | 0.40 |
|  | Pedal-a | 28 |  |  | 1.18 | 3.59 | 3.98 | 10.82 |
|  | Pedal-l-L | 29 |  |  | 0.32 | 0.93 | 0.94 | 0.98 |
|  | Pedal-l-R | 30 |  |  | 0.36 | 0.98 | 0.85 | 1.00 |
|  | Pedal-p | 31 |  |  | 1.43 | 4.28 | 5.20 | 11.03 |
| Magno complex | Magno-d-L | 32 |  |  | 0.38 | 1.09 | 1.52 | 2.95 |
|  | Magno-d-R | 33 |  |  | 0.38 | 1.19 | 1.62 | 2.77 |
|  | Magno-v-L | 34 |  |  | 0.12 | 0.33 | 0.30 | 0.29 |
|  | Magno-v-R | 35 |  |  | 0.12 | 0.31 | 0.27 | 0.36 |
|  | Magno-p-L | 36 |  |  | 0.41 | 0.53 | 0.91 | 1.24 |
|  | Magno-p-R | 37 |  |  | 0.45 | 0.50 | 0.85 | 1.21 |
| Palliovisc complex | PallioVis | 38 |  |  | 1.04 | 3.12 | 4.46 | 10.17 |
|  | PallioVis-lv-L | 39 |  |  | 0.51 | 0.34 | 0.88 | 1.21 |
|  | PallioVis-lv-R | 40 |  |  | 0.50 | 0.36 | 0.92 | 1.07 |
|  | FinL-L | 41 |  |  | 0.06 | 0.47 | 0.91 | 2.91 |
|  | FinL-R | 42 |  |  | 0.07 | 0.42 | 0.95 | 2.58 |
|  | Chrom-p-L | 43 |  |  | 0.19 | 0.14 | 0.81 | 1.72 |
|  | Chrom-p-R | 44 |  |  | 0.19 | 0.16 | 0.72 | 1.63 |
|  | Visc | 45 |  |  | 0.16 | 0.29 | 1.25 | 4.96 |
| Optic lobes | OPL-L | 46 |  |  | 43.63 | 157.50 | 188.64 | 334.56 |
|  | OPL-R | 47 |  |  | 46.81 | 155.50 | 184.37 | 356.52 |
|  | CC (mm^3) |  | 2.15 | 7.44 | 21.75 | 66.17 | 82.62 | 181.49 |
|  | OPLs (mm^3) |  | 7.11 | 28.82 | 90.44 | 313.00 | 373.01 | 691.08 |
|  | CNS (mm^3) |  | 9.26 | 36.26 | 112.19 | 379.17 | 455.63 | 872.57 |

**Table S3.** List of estimates of brain volume in cuttlefish and squid

| Species | ML(mm) | OPLs (mm<sup>3</sup>) | CC (mm<sup>3</sup>) | CNS(mm<sup>3</sup>) | VL (mm<sup>3</sup>) | OPLs/CNS(%) | VL/CNS(%) |
| --- | --- | --- | --- | --- | --- | --- | --- |
| *Sepia elegans* | - | 296.00 | 197.00 | 493.00 | 15.88 | 59.68 | 3.22 |
| *Sepia orbignyana* | - | 274.00 | 198.40 | 472.40 | 15.49 | 57.81 | 3.28 |
| *Sepia officinalis* (H) (Wild et al 2015) | 6.30 | 1.96 | 0.97 | 2.94 | n.a. | 66.85 | n.a. |
| *Sepia officinalis* (Maddock & Young 1987) | 80.00 | 232.40 | 163.80 | 396.20 | 1.19 | 58.66 | 0.30 |
| *Sepia officinalis* (Wirz, 1959) | - | - | - | - | - | 67.03 | 3.68 |
| *Sepia officinalis* (Frosch, 1971) | - | - | - | - | - | 70.73 | 2.03 |
| *Sepia bandensis* (Montague, et al 2022) | ca. 60 | 233.24 | 80.43 | 313.67 | 16.05 | 74.36 | 5.12 |
| *Sepia latimanus* (H) (Ziadi-Kunzli, et al 2019 ) | - | - | - | - | - | ca. 82 | - |
| *Sepia plangon* (H) | 8.00 | 7.11 | 2.15 | 9.26 | 0.22 | 76.78 | 2.38 |
| *Sepia plangon* (J) | 18.00 | 28.82 | 7.44 | 36.26 | 1.41 | 79.48 | 3.89 |
| *Sepia plangon* (J) | 32.00 | 90.44 | 21.75 | 112.19 | 4.14 | 80.61 | 3.69 |
| *Sepia plangon* (F) | 71.00 | 373.00 | 82.62 | 455.62 | 18.12 | 81.87 | 3.98 |
| *Sepia plangon* (M) | 73.00 | 313.00 | 73.30 | 386.30 | 15.51 | 81.03 | 4.02 |
| *Sepia plangon* (F) | 107.00 | 691.00 | 181.50 | 872.50 | 46.61 | 79.20 | 5.34 |
| *Sepioteuthis lessoniana* (Chung et al 2020) | 55.00 | 98.12 | 22.89 | 121.01 | 2.69 | 81.08 | 2.22 |
| *Sepioteuthis lessoniana* | 40.30 | 112.41 | 25.38 | 137.79 | 3.59 | 81.58 | 2.61 |
| *Sepioteuthis lessoniana* | 49.30 | 175.23 | 41.77 | 217.00 | 5.61 | 80.75 | 2.59 |
| *Sepioteuthis lessoniana* | 58.30 | 207.70 | 48.03 | 255.73 | 5.65 | 81.22 | 2.21 |
| *Sepioteuthis lessoniana* | 113.00 | 443.90 | 138.48 | 582.38 | 19.07 | 76.22 | 3.27 |

